# Shared neurophysiological resources between exogenous and endogenous visuospatial attentional processes

**DOI:** 10.1101/2022.04.11.487868

**Authors:** Mathieu Landry, Jason Da Silva Castanheira, Sylvain Baillet, Jérôme Sackur, Amir Raz

## Abstract

Prevailing accounts of visuospatial attention differentiate exogenous (involuntary shifts) from endogenous (voluntary control) orienting of attention. While these two forms of attentional processes are functionally separable, their interactions have been at the center of ongoing debates for more than two decades. One hypothesis is that exogenous and endogenous attention interfere because they share processing resources. Here, we confirm that endogenous attention alters exogenous attention processing, and examine the role of alpha-band neurophysiological activity in such interference events. We contrast the effects of exogenous attention across two experimental conditions: a single-cueing condition where exogenous attention is engaged alone, and a double-cueing condition where exogenous attention is concurrently engaged with endogenous attention. Our results show that the engagement of endogenous attention alters the emergence of exogenous attention across cue-related and target-related brain processes. Importantly, we also report that classifiers trained to decode exogenous attention from the power and phase of alpha-band brain activity in the single-cueing condition fail to do so in the double-cueing condition, where endogenous attention is also engaged. Taken together, our observations challenge the idea that exogenous attention operates independently from top-down processes and demonstrate that both forms of attention orienting engage shared brain processes, which constrain their interactions.

**Significance Statement:** Visuospatial attention is often dichotomized into top-down and bottom-up components: Top-down attention reflects slow voluntary shifts of attention orienting, while bottom-up attention is recruited by emerging demands from the environment. A large body of previous findings support the view that these two forms of attention orienting are functionally separable, with some interactions. The current study examines such interactions between top-down and bottom-up attention. Using electroencephalography (EEG) and multivariate pattern classification techniques, the researchers show that top-down attention interferes with the brain activity patterns of bottom-up attention. Moreover, machine learning classifiers trained to detect bottom-up attention based on brain activity in the alpha band (8-12 Hz), a marker of visuospatial attention, fail systematically when top-down attention is also engaged. The authors therefore conclude that both forms of visuospatial orienting are supported by overlapping processes that share brain resources.

## Introduction

The control of visuospatial attention is often dichotomized into exogenous and endogenous forms of attentional orienting. In this framework, exogenous orienting involves reflexive and involuntary shifts of attention triggered by salient events, while endogenous orienting corresponds to the voluntary control of attention based on behavioral goals (Corbetta et al., 2008; Corbetta & Shulman, 2002; Jonides, 1981; Müller & Rabbitt, 1989; Posner, 1980; Wright & Ward, 2008). A large body of work underlines critical differences between these two forms of attention orienting, in line with the idea that they map onto separate and independent brain systems (Chica et al., 2013). For example, exogenous orienting is induced by involuntary responses that quickly engage and disengage from attended locations in an automatic (i.e., occurring in the absence of voluntary control) and ballistic (i.e., unstoppable once engaged) fashion, whereas endogenous orienting reflects slower voluntary shifts of attention towards sensory events (Carrasco, 2011). In sum, visuospatial attention involves at least two systems that operate via distinct modes of control.

Studies of visuospatial attention have often used the spatial cueing experimental paradigm where exogenous orienting is induced using task-irrelevant and peripheral stimuli (e.g., a flash occurring at the periphery of the visual field) and endogenous attention is mobilized by cues instructing participants to voluntarily attend to a particular location (Chica et al., 2014). In this approach, the perceptual benefits of visuospatial attention are revealed by contrasting individual performances when target events occur at the cued versus uncued locations (Posner et al., 1980). Based on this paradigm, studies have examined whether exogenous and endogenous attention operate independently from each other using a double-cueing strategy where both forms of attention orienting are concurrently engaged. This double-cueing strategy reveals that exogenous and endogenous attention are not fully separable. Indeed, while some reports show limited behavioural interference between the two systems (i.e., both attention systems can independently improve performance on a perceptual task) (Landry et al., 2021), others have uncovered evidence of interferences both at the behavioral (e.g., Berger et al., 2005) and the neural (Busse et al., 2008; Hopfinger & West, 2006) levels. Furthermore, neuroimaging reports have also uncovered overlaps between exogenous and endogenous attention at the sensory levels (Dugué et al., 2020; Müller & Ebeling, 2008) and across brain networks (Bowling et al., 2020; Buschman & Miller, 2007; Nobre et al., 1997; Peelen et al., 2004). In sum, while exogenous and endogenous attention represent two distinct constructs of attention systems, they are not completely functionally independent from each other.

One hypothesis with regards to attention typologies and their interactions proposes that exogenous and endogenous attention processes represent separate functional systems capable of mutual interference at the perceptual level (Berger et al., 2005). Insofar that visuospatial attention facilitates perception by allocating processing resources at the sensory level (Luck et al., 1996), mutual interference between them would entail that both systems compete for the same limited perceptual resources (Busse et al., 2008). This hypothesis has several implications. First, it implies that exogenous attention, a ballistic and automatic process, is not impervious to the influence of endogenous attention. This hypothesis challenges the idea that exogenous attention operates independently from top-down processes (Pinto et al., 2013). Second, it also implies that the interactions between the limited neural processing resources shape the dynamics between exogenous and endogenous attention. Attention systems would therefore operate in accordance with the neural processing resources available, which means that when resources are strained and the two attention systems are engaged, we may predict that exogenous and endogenous attention interfere with one another.

In this regard, recent electrophysiology work proposes that exogenous and endogenous orienting both involve alpha-band activity as a common brain signal marker (8-12 Hz; Keefe & Störmer, 2021). This potential brain signal marker would enable the investigation of interactions between exogenous and endogenous attention using noninvasive neurophysiological methods. In the context of visuospatial attention, alpha-band signal power over the posterior cortex is viewed as a marker of cortical excitation and inhibition (Klimesch, 2012). Studies show that visuospatial attention links to decreased alpha-band activity in contralateral posterior visual regions, and increased alpha-band activity in ipsilateral visual regions (Peylo et al., 2021). This pattern accordingly implies that visuospatial attention operates by increasing neuronal excitation of contralateral sensory regions and increasing inhibition of ipsilateral sensory regions, suggesting a causal association between localized changes of alpha-band power and visuospatial attention behavior (Bagherzadeh et al., 2020). Furthermore, the phase of alpha oscillations also plays an important role for gating sensory inputs (Mathewson et al., 2009). Hence, both the power and phase of alpha rhythms reflect neural mechanisms of selective attention.

The purpose of the present study was twofold. First, we aimed to confirm that endogenous attention influences exogenous attention processing (Santangelo & Spence, 2008). Contrary to the idea that exogenous attention is impervious to top-down processes, we expected that endogenous attention would interfere with exogenous attention processing. Second, we planned to examine if this interference involves alpha-band activity. We hypothesized that the interactions between exogenous and endogenous attention would be captured by asymmetric changes of posterior cortical alpha-band power and phase. To test this hypothesis, we compared the behavioral and neurophysiological effects of exogenous attention in a single-cueing experimental condition, where exogenous attention is engaged alone, with respect to a double-cueing condition, where exogenous attention is concurrently engaged with endogenous attention. Our experimental approach was designed so that the endogenous cue always preceded the orientation of exogenous attention during double cueing. This means that modulations of endogenous attention over exogenous attention would reflect the involvement of top-down processes rather than a perceptual confound (i.e., changes in the visual display). We also separated the trials of the double-cueing condition into trials where both cues indicated the same location of the visual field (labelled as double cueing same location hereafter), and trials where they pointed to opposite locations (labelled as double cueing opposite locations hereafter). This experimental strategy enabled the study of situations where attention resources are distributed across the visual field.

To explore both hypotheses, we performed multivariate pattern classification on electroencephalography (EEG) signals in two separate decoding challenges. The first challenge tested whether classifiers trained to decode the cue-related and target-related effects of exogenous attention in the single cueing condition would generalize to the double-cueing condition. Here, we expected that the classifiers’ performances would decrease if endogenous attention interferes with exogenous attention processes. The second challenge tested whether we could decode the cueing condition (i.e., single versus double-cueing) from EEG data. Above chance decoding of the cueing condition would therefore be indicative of differences in exogenous orienting between the single and double cueing conditions, and therefore demonstrate that endogenous attention interferes with exogenous orienting. Both decoding challenges complement each other: The first challenge examines the performance of classifiers trained only to decode the cue-related and target-related effects of exogenous attention in a new context involving endogenous attention; the second challenge focuses directly on the cueing condition. This distinction is important because the first challenge validates that one can decode the effects of exogenous attention, whereas the second one directly assesses the decoding of the cueing condition (i.e., single versus double cueing).

## Methods

### Participants

We recruited 42 participants (29 women, mean age = 22.2 (sd = 3.09)) based on convenience sampling. Each participant received monetary compensation for completing 4 different tasks during one testing session lasting approximately 2 hours. All participants had normal or corrected-to-normal vision, had no prior history of neurological disease, and provided consent. The experiments were approved by the McGill Faculty of Medicine & Health Sciences Research Ethics Board (Certificate #511-0516).

We excluded data from ten participants: Seven participants were excluded due to poor EEG data quality. One participant did not complete all conditions. We excluded another participant because they performed below chance-level (~40% averaged across all conditions). Lastly, one participant was excluded due to a high volume of timeout errors (response times > 1500ms on ~11% of trials overall). Data analysis therefore proceeded from the data of 32 individuals (24 women, Mean age = 21.9 (SD = 2.6)). We determined our sample size based on previous work using the double-cueing experimental approach (Landry et al., 2021). Simulations of the effect size determined that 6 participants were required to achieve a power of .8 for the within-individual cueing effects of exogenous and endogenous orienting in a target discrimination task, wiht an alpha set at 0.05. The present sample size was therefore adequate to detect the desired attention effect at the behavioral level, and about twice that of previous EEG studies of combined effects of exogenous and endogenous orienting (Hopfinger & West, 2006; Keefe & Störmer, 2021).

### Stimuli, Apparatus & Design

Participants viewed tasks on a 24-in BenQ G2420HD monitor positioned approximately 60 cm in front of them. Stimulus presentation was controlled via MATLAB R2015b (Mathworks Inc., Natick, MA, USA) using the third version of the Psychophysics toolbox (Brainard & Vision, 1997; Kleiner et al., 2007; Pelli, 1997). The screen refresh rate was set to 75 Hz. Stimuli consisted of black (i.e., RGB values of 0, 0, 0; 1.11 cd/m^2^) and white (i.e., RGB values of 255, 255, 255; 218.8 cd/m^2^) drawings on a grey background (i.e., RGB values of 128, 128, 128; 70.88 cd/m^2^). The fixation marker consisted of a black circle subtending a radius of 1.2° located in the center of the screen. Two target placeholders were located 8.7° away from the fixation marker on the left and right side of the screen. These placeholders consisted of black circles with a 2.4° radius. We cued exogenous attention by briefly changing the line drawing from one of the placeholders to white. To ensure that this cue solely engaged exogenous orienting, the cue-target spatial contingency was set to 50%, such that the cue was task-irrelevant and not predictive of the target’s location per the standard in the field (Chica et al., 2014). We cued endogenous attention by coloring the inside of the fixation marker, wherein the right or left half of the circle was shaded in black and the other half in white. The side of the fixation marker that turned white indicated where the target was likely to occur. For example, if the right side turned white, the target was 66.6% likely to appear in the right placeholder. In this way, we avoided using overlearned directional cues, such as when using an arrow, to engage endogenous attention (Ristic & Kingstone, 2012; Ristic & Landry, 2015). The targets were sinusoidal black and white gratings modulated by a Gaussian envelope with a spatial frequency of 3 cpd and tilted 5° clockwise or counterclockwise from the vertical.

### Procedure

Participants completed four conditions: A non-cueing condition, a single exogenous cueing condition, a single endogenous cueing condition, and a double-cueing condition that involved both exogenous and endogenous orienting. Participants performed 10 practice trials before each condition. Each condition consisted of 384 trials each. The order of conditions was randomized across participants. Here we present results from the non-cueing, the exogenous cueing, and the double-cueing conditions.

Participants were instructed to fixate the center of the screen throughout the experiment, while we monitored their eye movements using electro-oculogram. We jittered the latencies between the respective onsets of all events in the tasks i.e., fixation point, attention orienting cues, and the target event, based on a uniform distribution (range detailed below), to minimize the effects of temporal predictability following spatial cueing. The latencies between attentional cues and the target were adjusted based on previous literature of exogenous and endogenous attention, see description of the tasks below (Chica et al., 2014). We chose these latencies (detailed below and in Figure 1) such that each attention system would have maximal behavioural facilitation.

**Figure 1.**
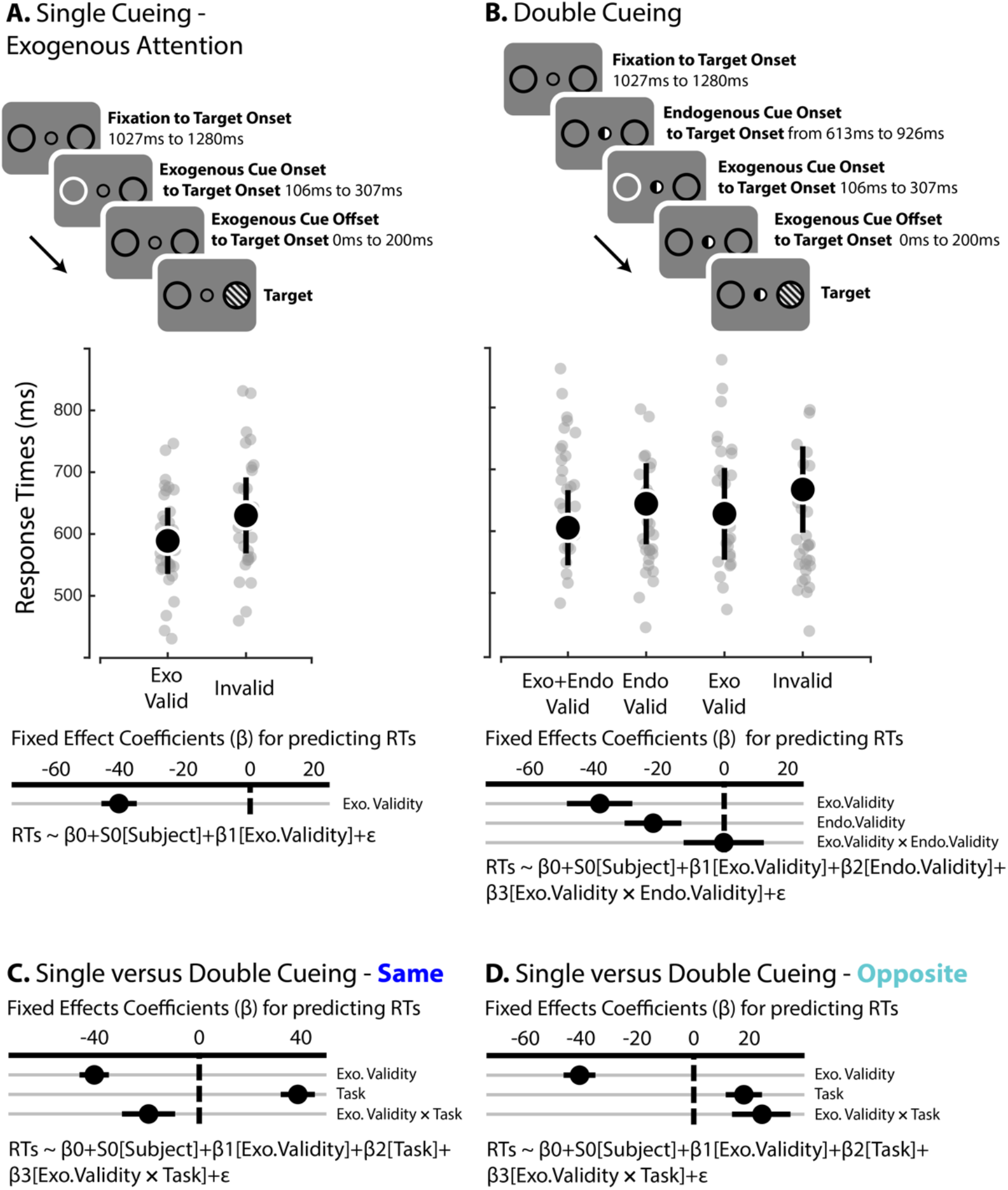
**A.** Top panel: in the single-cueing condition each trial began with a fixation screen, followed by the onset of the exogenous cue and lastly, the Gabor target at the valid or invalid location. The exogenous cue was task-irrelevant and predicted the upcoming target’s location at chance-level. The scatter plot depicts group (black dots) and participant (grey dots) average RTs for each condition. Error bars represent bootstrapped 95% C.I. Bottom panel: single-trial hierarchical regression model coefficients. Error bars represent 95% C.I. **B.** Top panel: the double-cueing condition comprised both exogenous and endogenous cueing where the exogenous cue did not predict the target’s location and the endogenous cue predicted the target’s location 66.66% of the time. The endogenous cue (i.e., fixation semicircles, where the white side indicated where the target was likely to occur; see Methods for details) appeared after a jittered interval, followed by the exogenous cue, and finally the target event. The scatter plot shows group (black dots) and subject (grey dots) average RTs. Error bars represent bootstrapped 95% C.I. Bottom panel: single-trial hierarchical regression model coefficients. Error bars represent 95% C.I. **C.** Single-trial hierarchical regression model coefficients where evaluated the effect of cueing condition on exogenous cue validity.

In the non-cueing condition, the fixation stimulus was displayed over 1027 ms to 1280 ms, and was immediately followed by the onset of the target event, which remained on the screen until the participants’ response. The non-cueing condition was used as a baseline condition against which we regressed the sensory effects of cue stimuli for the target-related analyses (see the *Electroencephalography* section below).

We used the same jittered timing interval between the fixation and the target event for the single exogenous cueing condition. We similarly varied the onset of the exogenous cue from 106 ms to 307 ms before the target event. For all trials, the exogenous cue remained on the screen for 106 ms. The target remained on the screen until the participants’ response.

We used the same timing for the double-cueing condition between fixation and the target event. Participants were first presented with the endogenous cue which preceded the target stimulus from 613 ms to 926 ms. The endogenous cue remained on the screen until the end of the trial. Again, the exogenous cue would then be displayed from 106 ms to 307 ms before target onset, and remained on the screen for 106 ms (Figure 1).

As mentioned above, the exogenous cue was uninformative about the target’s location, whereas the endogenous cue was made task-relevant by indicating the correct target location in 66.6% of the trials. Participants were aware of these contingencies. They were instructed to discriminate the orientation of the Gabor target as quickly and accurately as possible by pressing the F key of a QWERTY keyboard to indicate counterclockwise orientation, and the J key to indicate clockwise orientation. The inter-trial period was set to 1s.

### Behavioral performances

Participants’ discrimination performances were near ceiling: the average accuracy rate was about 93%. We examined response times (RTs) for accurate trials only. We discarded trials where participants anticipated the target onset (i.e., RTs < 150ms) or did not reply (i.e., RTs > 1500ms), which accounted for less than 2% of the trials, as well as trials with wrong key presses (neither F nor J, <1% of all trials). We used hierarchical linear regression models (Gelman & Hill, 2006) to test the effects of cueing (i.e., valid versus invalid) and condition (i.e., single versus double cueing), wherein predictors were added in a stepwise fashion. We applied a chi-square goodness-of-fit test over the deviance to determine whether predictors significantly improved the fit, and assessed Bayesian information criterion (BIC) to select the best fitting model. We also estimated Bayes factor weighting evidence for the null against the alternative hypothesis for one degree of freedom (Wagenmakers, 2007):

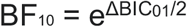

Where the delta BIC is the difference between the Bayes Criterion Information for the null model and the Bayes Criterion Information for the alternative model.

We ran four statistical analyses on the behavioural performances of participants. First, we ran a linear regression analysis of the single exogenous cueing condition with cue validity as a fixed factor. Second, for the double-cueing condition, we ran a linear regression analysis where exogenous cue validity, endogenous cue validity, and their interaction were fixed predictors. Because we focused on comparing the single and double-cueing conditions, we examined if the cueing effect of exogenous attention varied as a function of cueing condition (i.e., single and double cueing). We accordingly ran a linear regression analysis where exogenous cue validity and cueing conditions (i.e., single versus double-cueing conditions), as well as their interactions, were fixed factors. We did so separately for trials where both cues indicated the same location, and for trials where both cues indicated opposite locations. We included participants as a random factor for all regression models.

### Electroencephalography

We recorded the EEG using 64 Ag/AgCl active electrodes (ActiCap System; Brain Products GmbH; Gilching, Germany) positioned following the 10-20 system, at a sampling rate of 1000 Hz. We also monitored eye blinks and eye movements through bipolar electrodes placed at the outer canthi, as well as the superior and inferior orbits of the left eye. Impedances of all electrodes were kept below 10 kΩ. All electrophysiological signals were amplified (ActiChamp System; Brain Products GmbH; Gilching, Germany). Electrodes were referenced online with respect to C4, and rereferenced offline to an average reference. Preprocessing and analyses were conducted with BrainVision Analyzer (ActiChamp System; Brain Products GmbH Inc.; Gilching, Germany), Brainstorm (Tadel et al., 2011) and custom Matlab (R2020a; Mathworks Inc., Natick, MA) scripts. We downsampled the data to 250 Hz and applied two IIR Butterworth filters: a high-pass filter of 8th order above 0.1 Hz and a 60-Hz notch filter. We then topographically interpolated bad channels (1.12% of channels across participants). Next, we performed an independent component analysis to identify artifacts caused by eye movements and blinks using the BrainVision Analyzer Ocular correction ICA tool. We corroborated these analyses with the actual electro-oculographic traces. We corrected the EEG signals accordingly by remixing the artifact-free independent components together. We visually inspected the EEG traces further to identify segments affected by atypical noise patterns, as well as portions where EEG amplitude exceeded ± 200 μV across all electrodes. All event triggers were temporally realigned based on photodiode recordings of the actual onsets of the exogenous cues, endogenous cues, and target events.

Given the temporal proximity of cueing and target events, neurophysiological activity due to the cues overlapped with that of the target stimulus. Hence, to minimize the contamination of the target-related ERP from the cue events, we subtracted the electrographic response of the exogenous and endogenous cues presentations whenever we examined target-related response. However, in lieu of the adjacent response filter method often used for this purpose (Woldorff, 1993), we removed cue event-related responses by taking the residuals of the linear regression model using the non-cueing condition as baseline. We accordingly ran a hierarchical linear regression analysis with condition as a dummy coded fixed factor, and subjects as a random factor. Raw residuals were obtained from the Matlab function *residuals()* for all channels and subjects.

### Event-related potentials

We obtained cue-related and target-related event-related potentials by averaging EEG epochs across trials and participants. We applied a FIR bandpass-pass filter between 0.5 and 15 Hz and epoched the EEG recordings around - 200 and 1000 ms with respect to cue or target events. The resulting ERPs were baseline corrected (−100 to 0 ms).

### Lateralized alpha index

We computed the lateralized alpha index (LAI) to characterize the time-resolved changes of alpha power with experimental conditions. We first estimated the instantaneous power of alpha-band EEG activity over the entire epoch duration (from −200 to 1000 ms related to the exogenous cue onset) using the squared modulus of the Hilbert transformation of the EEG traces, and then computed the LAI following the direction of the cue at each time point according to the following equation:

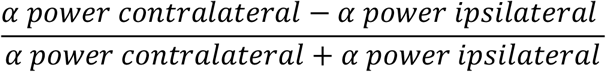

The contra- and ipsilateral sides were defined with respect to the location of the exogenous cue. We computed the LAI for all lateral EEG channels (i.e., excluding all midline ‘z’ channels), and paired every lateral channel with the homologous opposite channel (e.g., O1 and O2). We then standardized the LAI via a z-score transformation with respect to the mean and standard deviation of the LAI from their baseline statistics (−100 to 0 ms).

### Intertrial phase clustering

To examine the phase component of alpha oscillations, we computed the inter-trial phase clustering (ITPC) measure for exogenous cue-related EEG activity across all channels between −200 and 1000ms with respect to cue onset (Tallon-Baudry et al., 1996) The ITPC derivation uses the imaginary part of the Hilbert transform to estimate the instantaneous phase angle of the ongoing alpha cycles (Cohen, 2014):

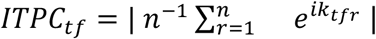

Where, *n* corresponds to the number of trials and *e^ik_tfr_^* is the complex polar representation of the phase angle *k* on trial r for the index for time and frequency tf. Lastly, we z-scored transformed the ITPC indices from their baseline statistics (−100 to 0 ms).

### Multivariate decoding analyses

We evaluated whether endogenous attention impacts the neurophysiological markers of exogenous attention using a multivariate pattern classification approach (Hong et al., 2020). We used a decoding approach developed by Bae and Luck (2018).

We trained a linear support vector machine (SVM) classifier at each time point using MATLAB’s *fitcsvm* function. We randomly split trials of each participant into three equally sized bins and averaged the neurophysiological features within each bin (see *Challenge I: Generalization of decoding going from single to double cueing and Challenge II: Decoding cueing conditions* for a description of the features used to train the classifiers). Note that the bin size varied between cue-related and target-related analyses (see below for exact bin size in each analysis separately). This produced three distinct time-series data to train the classifier. We then used a three-fold cross-validation procedure, whereby 2 of the 3 bins were used to train the classifier, and the remaining bin was used to test the SVM performances. Classifiers were trained and validated at each timepoint for every participant across the EEG time series (see below for details). Decoding accuracy was established on correct classification at each time point when we applied the trained SVM model to the validation set. We iterated across all possible combinations (3 possibilities) of training and testing sets based on the three averaged time-series: Each averaged time-series was used twice for training the SVM classifiers and was used once to validate an SVM classifier. We repeated the above random binning, averaging, and decoding procedure 50 times per participant (see supplementary Figure 1).

We averaged decoding accuracies across participants, for each cueing condition, and neurophysiological features. We applied a five-sample sliding window, i.e., a 20-ms window, to smooth the average decoding accuracy time series. We used cluster-corrected permutation tests to control for family-wise type-II error rates. We summed t-statistics from adjacent significant time points (p < .05), which provided a mass t-value for each cluster. We then performed 1000 permutations where we randomly shuffled the class labels for each time series and then computed mass t-statistics for each permutation. We then estimated the p-value for the observed cluster by comparing the mass t-statistics of each cluster against surrogate distributions for the null hypothesis. Significant cluster size was set to p<.05. We applied this statistical approach for all decoding analyses across two separate decoding challenges (see supplementary Figure 1). First, we evaluated the ability of classifiers trained on the single-cueing condition to generalize to the double-cueing condition (i.e., decoding Challenge I). Second, we trained linear SVM classifiers to decode the cueing condition, i.e., single versus double cueing (decoding Challenge II). This complementary analysis aimed to corroborate our findings from Challenge I where we show that classifiers trained in the single cueing condition performed worst in the double-cueing condition. Hence, Challenge II was designed to confirm that cue-related and target-related effects of exogenous attention differ between cueing conditions.

### Challenge I: Generalization of decoding going from single to double cueing

We assessed how classifiers trained in the single cueing condition generalized to the double cueing condition. We rationalized that a decrease in decoding performance from the single-cueing to the double-cueing condition would reflect interference of endogenous attention over exogenous attention, i.e., that the effects of exogenous attention are not impervious to the concurrent engagement of endogenous attention (see supplementary Figure 1b). For the double cueing condition, we assessed trials for same and opposite cueing locations separately consistent with our hypothesis that cueing opposite locations would put greater strain on attention resources.

We applied this decoding approach to both target-related and cue-related neurophysiological features of exogenous attention. This analysis tested whether the processing of the exogenous cue and the target stimulus were altered by endogenous attention processes.

We trained classifiers to determine the direction of exogenous attention orienting (left versus right orienting) to examine cue-related effects and exogenous cue validity (exogenous cue valid versus invalid) for target-related effects. We first decoded cue-related and target-related exogenous attention based on ERP neurophysiological signals. For cue-related activity, each bin used to produce averaged time series for decoding comprised 25 trials. For target-related activity, each bin used to generate the averaged time series for decoding included 21 trials. Furthermore, to ensure that decoders were not influenced by the target’s location during target-related decoding, we trained them separately for left and right target locations, before averaging their respective classification performances (i.e., averaged decoding accuracy of right and left target location).

To test our main hypothesis about alpha-band activity, we then decoded the direction of exogenous orienting from cue-related LAI and IPTC patterns. Each bin used to produce averaged time series for decoding comprised 25 trials.

For Challenge I, we first computed t-statistics at each time point to assess decoding performances against chance-level (i.e., 50% decoding accuracy) to validate the SVM approach. Next, we used pairwise t-statistics to compare: i) single-cueing against double-cueing decoding performances, when both cues indicate the same direction; ii) single-cueing against double-cueing, when both cues indicate opposite directions, and iii) double-cueing with both cues indicate the same direction vs. opposite directions.

### Applying principal component analysis to the processing of the exogenous cue

We aimed to isolate which components of the exogenous cue processing were impacted by endogenous attention. To this end, we examined the encoding model of the classifiers. We trained SVM classifiers to decode the direction of exogenous attention based on the three-fold cross-validation approach (see *Multivariate decoding analyses*.). Training and validation were completed separately for single exogenous cueing, double cueing same location, and double cueing opposite locations. We therefore divided trials into separate bins for all three conditions – i.e., 16 trials per bin. We evaluated the decoding performances against chance levels for all conditions. Next, we examined the encoding model by extracting beta weights from each classifier at each time point and multiplied them by the covariance of the corresponding ERP data (Haufe et al., 2014). We averaged these time series of SVM beta weights across all participants for each condition separately and applied principal component analysis (PCA) to the averaged SVM beta weights from single cueing, double cueing same location, double cueing opposite locations. Next, we rotated the loadings using varimax rotation and kept the first two components, which explained 84.5% of the total variance. We projected the averaged SVM weights from each participant and cueing conditions onto both dimensions. Lastly, we compared all three cueing conditions using cluster-corrected one-sample t-tests across the time series of SVM weights and PCA components separately.

### Challenge II: Decoding cueing conditions

We next aimed to decode cueing conditions (i.e., single cueing versus double cueing). This analysis follows from the first decoding challenge. In Challenge I, we tested if decoding of exogenous attention generalizes across contexts with the hypothesis that classifiers trained to decode exogenous attention will fail to generalize to the double cueing condition due to interference from endogenous attention. With Challenge II, we aim to clarify a related question: can a decoder classify whether exogenous orienting was engaged by itself vs with endogenous attention (i.e., single vs double cueing conditions). If endogenous attention does not alter exogenous orienting, then a classifier trained to decode cueing conditions should be at chance-level. We hypothesize that we can decode cueing conditions above chance level due to the interference of endogenous attention on exogenous orienting. We followed a similar three-fold procedure as described previously, except for the following changes described below.

For exogenous cue-related EEG, we examined the cue-locked ERPs, cue-locked LAI, and cue-locked lateralized alpha ITPC. We averaged trials from bins separately for ipsilateral and contralateral channels and then subtracted the resulting time series from one another (i.e., contralateral minus ipsilateral). This process resulted in three averaged time series that we used for decoding, akin to the procedure above. Each bin contained 32 trials. Again, we separated trials in the double cueing condition into trials where both cues pointed to the same direction and trials where cues pointed in opposite directions

For target-related EEG, we used the exogenous cue-validity ERPs. We averaged trial bins separately for cue valid and invalid trials and then subtracted these waveforms (i.e., exogenous cue valid minus exogenous cue invalid). Each bin used to compute the average time series contained 14 trials. We trained and validated our SVM classifiers on these exogenous cue validity waveforms. We performed this analysis across contralateral and ipsilateral channels separately and split trials in the double cueing condition into trials where the cues indicated to the same or opposite locations.

We also decoded the cueing conditions (i.e., single versus double cueing) based on exogenous cue validity for target-related LAI. This analysis solely involved comparing single cueing to double cueing where cues indicated opposite locations. Again, we averaged trial bins separately for cue valid and invalid trials and then subtracted these averaged patterns (i.e., exogenous cue valid minus exogenous cue invalid). SVM classifiers were trained and validated on these averaged patterns. Each bin contained 16 trials. Furthermore, we extracted the beta weights of the classifiers across the time series using the analytic approach described by Haufe et al. (Haufe et al., 2014) to assess the topographical pattern associated with classification performances.

## Results

### Behavior

For the single cueing condition, we found that exogenous cueing improved discrimination performances, wherein cue validity was a reliable predictor (β=−41.28, SE=2.83, 95% CI [−46.83, −35.74]; Figure 1A and Supplementary Tables 1 and 2). In the double cueing condition, our behavioral analysis shows that the main effects of exogenous cue validity (β=−39.27, SE=3.04, 95% CI [−45.22, −33.32]) and endogenous cue validity (β=−22.42, SE=3.22, 95% CI [−28.73, −16.11]; Figure 1B, and Supplementary Tables 3 and 4) were statistically also reliable. This outcome confirms the engagement of both exogenous and endogenous attention in the double-cueing condition. Moreover, we observed that evidence weighted against their interaction, as indicated by a Bayes factor analysis: BF_01_=105.59).

Next, we examined whether exogenous cue validity varied between the single and double-cueing conditions. When both cues indicate the same location, the best-fitting model included exogenous cue validity (β=−41.28, SE=2.93, 95% CI [−47.02, −35.55]), cueing condition (β=39.98, SE=4.18, 95% CI [31.78, 48.18]), and their interaction (β=−19.91, SE=5.31, 95% CI [30.33, −9.49]) as predictors of RTs (Figure 1C, and Supplementary Tables 5 and 6). We then examined the cueing effect of exogenous attention between the single and double-cueing conditions when the cues indicated opposite locations. The best-fitting model included exogenous cue validity (β=−41.31 SE=2.93, 95% CI [−47.02, −35.55]), cueing condition (β=18.13, SE = 3.34, 95% CI [11.59, 24.69]), and their interaction (β=24.48, SE = 5.37, 95% CI [13.98, 35.07]) as predictors of RTs (Figure 1D, and Supplementary Tables 7 and 8). Together, these results show that concurrent endogenous attentional cueing alters exogenous cueing effects.

### Electrophysiology. Challenge I: Generalization of decoding going from single to double cueing

We first validated our decoding approach by showing that classification was above chance level for decoding the location of the exogenous cue across single cueing and double cueing conditions (supplementary Figure 2A). This result validates that our SVM approach can accurately decode the location of the exogenous cue during single cueing, and that this decoding ability transfers from single to the double cueing condition.

We then compared classification performances across single cueing, double cueing for the same location, and double cueing for opposite locations. These contrasts reveal the impact of endogenous attention on the processing of the peripheral cue. Classification performance was worse in the double-cueing condition compared to the single condition for exogenous cue-related ERPs (Figure 2A). We found significant differences in decoding accuracy when contrasting single cueing against double cueing for the same location between 144 ms and 496 ms. In turn, we found an early cluster (i.e., from 108 ms to 180 ms) and two late clusters (i.e., from ~448 ms to ~596 ms and from 636 ms to 704 ms) when we compared single cueing and double cueing opposite locations. These results are consistent with our behavioral data and confirm that the concurrent engagement of endogenous attention modulates the processing of the peripheral cue, both when exogenous and endogenous attention point at the same or opposite locations. Furthermore, we observed a late significant cluster from 628 ms to 740 ms when contrasting trials where the cues indicate the same direction against trials where the cues indicate opposite locations in the double cueing condition. These results are also consistent with our behavioral findings of endogenous attention modulating the effects of exogenous attention.

**Figure 2.**
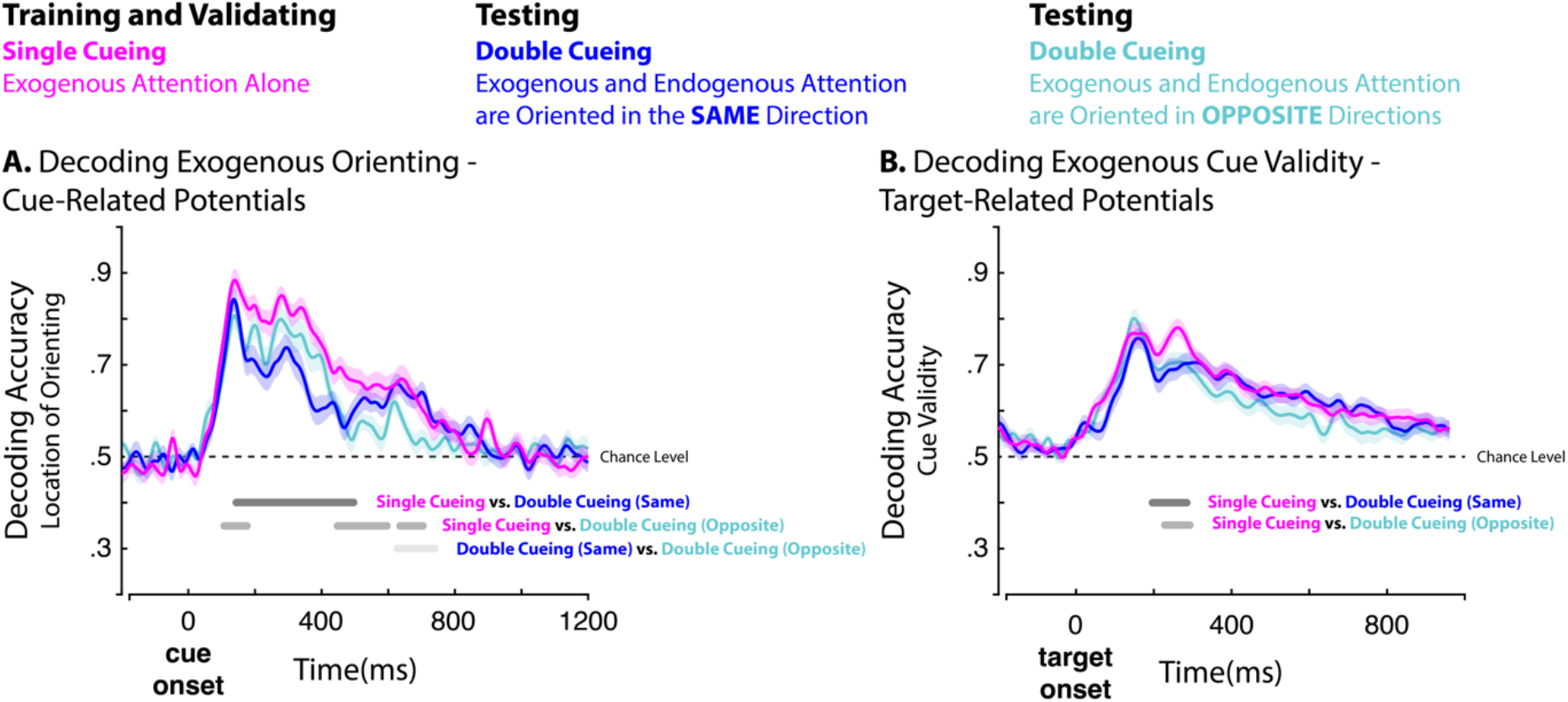
Multivariate classification of exogenous orienting and cue validity in the single and double-cueing conditions. We separately examined trials in the double-cueing condition where the cues indicated the same or opposite locations. We compared classification performances using cluster-corrected pairwise t-tests. Grey lines indicate significant clusters for each comparison. Shaded areas represent the SEM. **A.** Decoding accuracy of the direction of exogenous orienting based on cue-related ERPs. **B.** Decoding accuracy of exogenous attention cue-validity based on target-related ERPs.

We further investigated whether endogenous attention impacts the cueing benefits of exogenous attention at the neural level by decoding exogenous cue validity based on target-related waveforms. We first validated that classification was above chance-level in the single- and double-cueing conditions (supplementary Figure 2B), and that SVM could decode exogenous cue validity in the double-cueing condition (supplementary Figure 2B). Next, we compared our ability to decode cue validity across the three cueing conditions (i.e., single cueing, double cueing for the same location, and double cueing for opposite locations). Our decoding results were consistent with our behavioral findings, showing that classification performance for exogenous cue validity decreases for the double cueing compared to single cueing (Figure 2B). We observed significant differences between 196 ms and 284 ms for single vs double cueing same location, and between 228 ms and 292 ms for single vs double cueing opposite locations.

We then tested our main hypothesis by decoding the direction of exogenous attention based on alpha-band dynamics. The validation procedure first confirmed that classification was above chance level for both the LAI and ITPC features in the single cueing condition, thereby confirming that both features of alpha waves–i.e., LAI and ITPC–uncovered the effects of exogenous attention (supplementary Figure 3A and 3B). Next, we tested how these classifiers performed in the context of double cueing. Here, we observed a similar pattern for LAI and ITPC: Classification performance was above chance level when the cues indicated the same location, and below chance level when they indicated opposite locations (Figure 3; supplementary Figure 3A and 3B). We found significant clusters between 72-368 ms using LAI features and from 0-496 ms using ITPC features when we compared decoding accuracies between double-cueing same location and double-cueing opposite locations (Figure 3). In sum, classifiers trained to decode the direction of exogenous attention are confused by the orientation direction of endogenous attention. This outcome provides compelling evidence for the notion that both attention processes modulate lateralized alpha-band power and phase.

**Figure 3.**
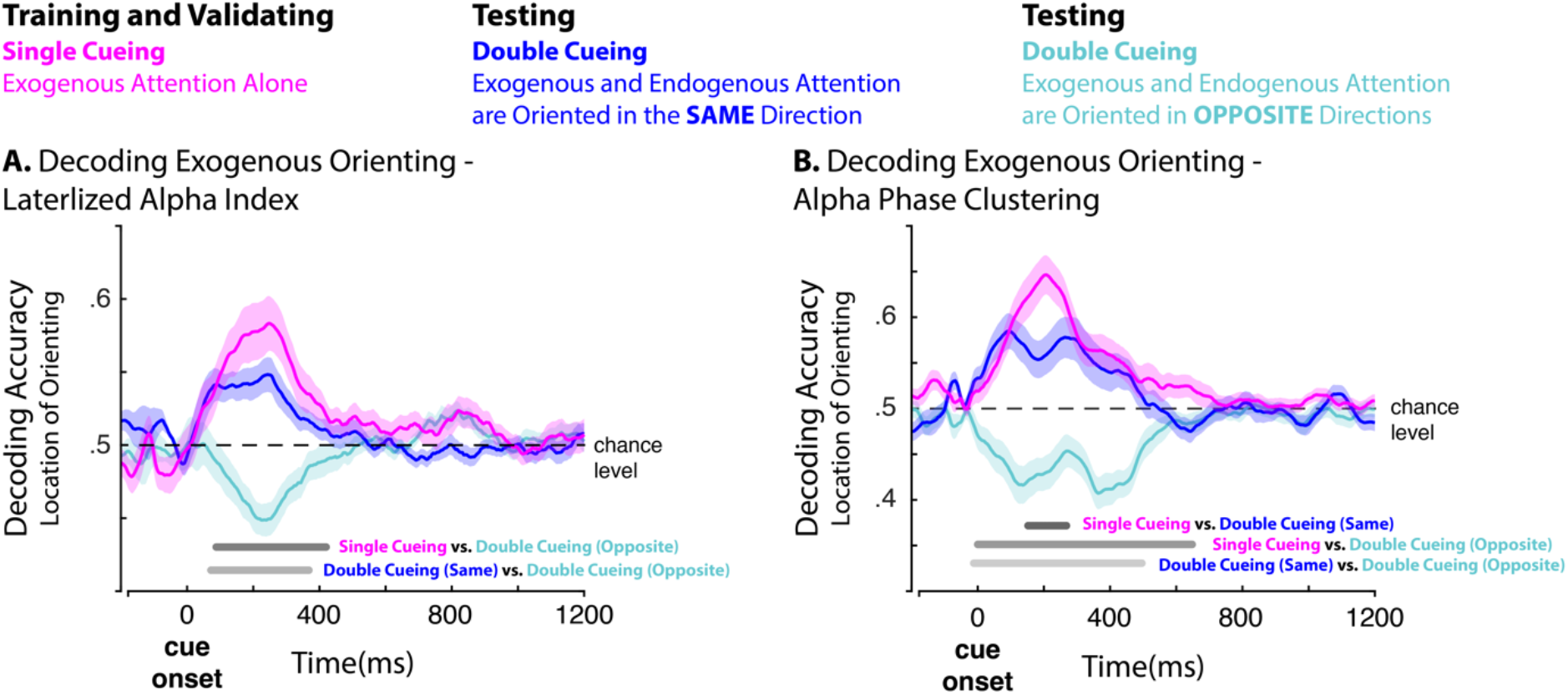
Multivariate classification of exogenous orienting and cue validity in the single and double cueing conditions. We examined separately trials in the double-cueing condition where the cues indicated the same or opposite locations. We compared classification performances using cluster-corrected pairwise t-tests. Grey lines indicate significant clusters for each comparison. Shaded areas represent the SEM. **A.** Decoding accuracy of exogenous cue orienting based on cue-related LAI features (see Methods). **B.** Decoding accuracy of exogenous orienting based on cue-related alpha ITPC features.

### Electrophysiology

#### Principal component analysis of SVM weights of exogenous attention

Since the decoding of exogenous attention did not generalize to the double cueing condition, we aimed to quantify what aspects of the cue-related ERPs were different between cue conditions. To do this, we trained classifiers on cue-related ERPs in the single and double cueing conditions separately (see supplementary Figure 4 for the averaged classification accuracy). We then extracted their SVM weights for each of the three classifiers (single, double-cueing same location, and double-cueing opposite locations) based on the Haufe et al. procedure (Haufe et al., 2014). We applied principal component analysis (PCA) separately to the SVM weight time-series of each classifier to reduce the dimensionality of the data and identify the relevant components. This produced two significant components.

The first component (57% of explained variance) was characterized by a posterior topography and early activity (> 200 ms) following cue onset (see supplementary Figure 2 and Figure 4A). We compared the three classifiers along the time-series of this early posterior component. We observed two significant clusters when comparing single cueing to double cueing opposite locations: A first one ranging from 204 ms to 268 ms, and a second one from 360 ms to 456 ms (Figure 4A). We also found two significant clusters when we compared double cueing same location versus double cueing opposite locations: A first one from 156 ms to 472 ms, and a second one from 560 ms to 668 ms (Figure 4A).

**Figure 4.**
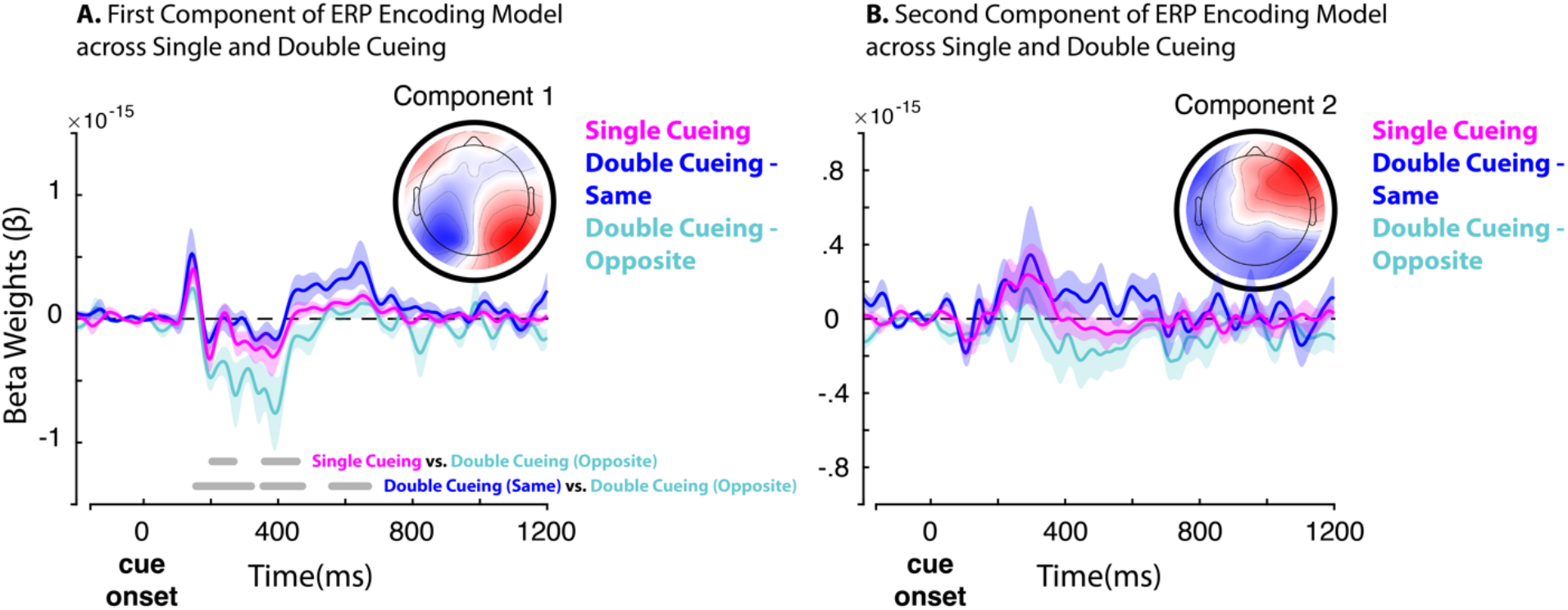
Comparison of single (pink), double cueing for same cueing location (blue), and double cueing for opposite cueing locations across first (**A**) and second (**B**) PCA components for the exogenous cue-related encoding model. Grey lines indicate significant clusters based on cluster-corrected pairwise t-tests. Bold lines represent the averaged time series and shaded areas of the SEM.

The second component from our PCA analysis (29% of explained variance) was characterized by a diffuse frontal topography and later activity (< 200 ms; see supplementary Figure 2 and Figure 4B). We, however, did not observe significant differences between the cueing conditions for the second component (Figure 4B).

#### *Electrophysiology*. Challenge II – *Decoding cueing conditions*

We subsequently aimed to decode cueing conditions (i.e., single cueing versus double cueing). Permutation tests show that SVM classifiers performed better than chance level at decoding the cueing conditions based on cue-related waveforms (Figure 5A and 5D). We observed significant clusters ranging from 140 ms to 476 ms when both cues indicated the same location for double cueing, and from 108 ms to 614 ms when both cues indicated opposite locations for double cueing. We similarly decoded cueing condition from alpha-band (8-12 Hz) power and phase. We observed significant clusters for LAI and ITPC only when both cues indicated opposite locations during double cueing (Figure 5B, 5C, 5E, 5F). This result aligns with the decoding results of Challenge I (Figure 3). Here, we observed a significant cluster between from 88 ms to 432 ms after cue onset for LAI (Figure 5E), and between −24 ms to 492 ms after cue onset for ITPC (Figure 5F). These results suggest that alpha-band features enable the decoding of exogenous attention when it is deployed in isolation or in combination with endogenous attention.

**Figure 5.**
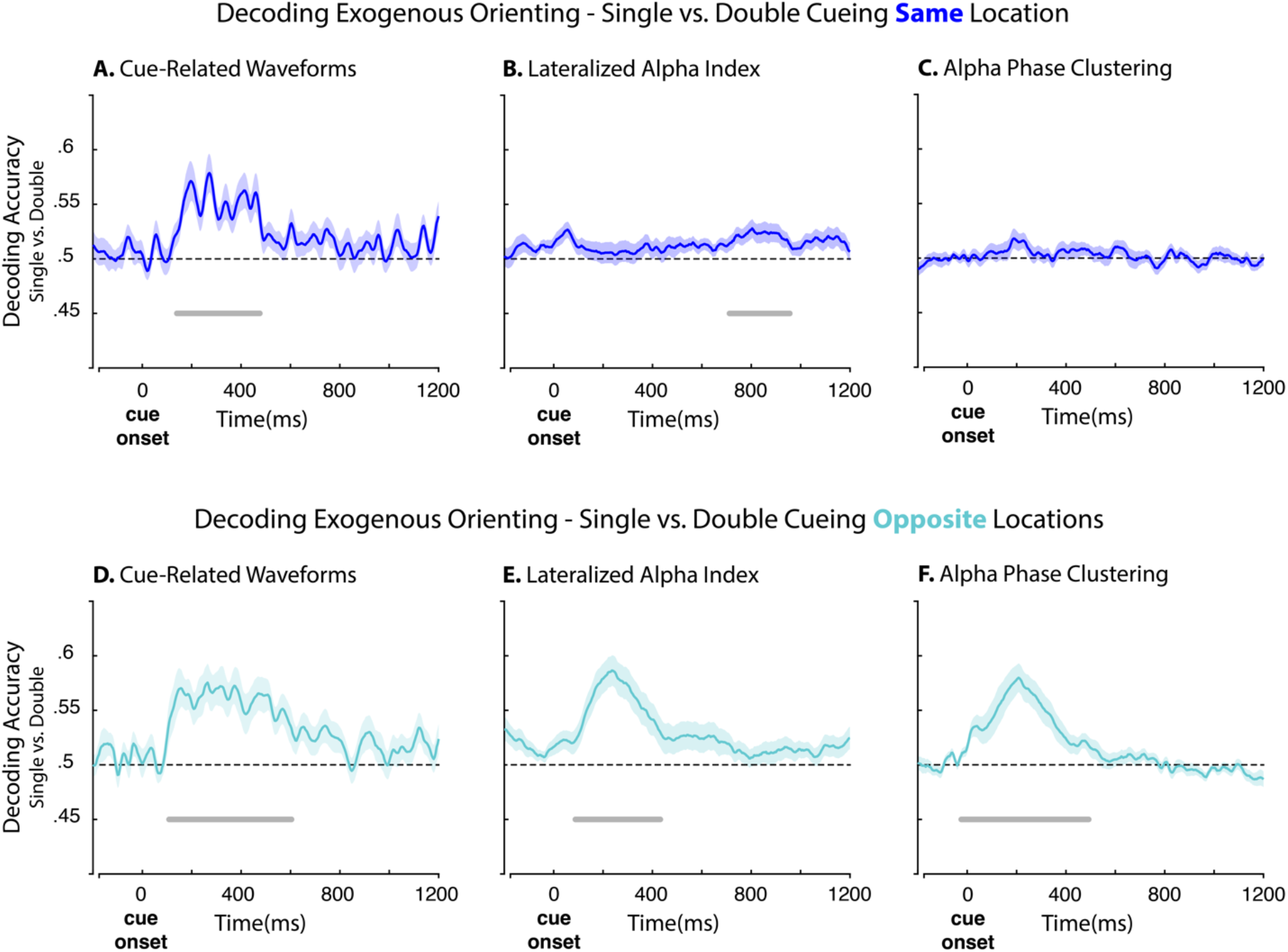
Multivariate classification of cueing conditions based on cue-related ERPs, LAI, and phase clustering. We assessed classification performance separately for trials in the double cueing condition where both cues indicated the same location (**A,B,C)** and opposite location (**D,E,F)**. Grey lines indicate above chance-level classification based on cluster-corrected t-tests. Bold lines represent the averaged time series and shaded areas the SEM.

We also classified cueing conditions from target-related ERPs. We found that classification performance was above chance level for both contra- and ipsi-lateral trials (Figure 6) regardless of cue location (i.e., same location vs opposite). We observed a significant cluster between 236 ms and 304 ms for the contralateral side (Figure 6A), and a cluster between 72 ms and 372 ms for the ipsilateral side (Figure 6B) when the cues pointed to the same direction (i.e., double-ceiling same location). In contrast, we observed a significant cluster only for ipsilateral trials between 168 ms and 280 ms when the cues pointed to opposite locations (Figure 6C, 6D). These results confirm the influence of endogenous attention over the cueing benefits of exogenous attention, even when both attention processes are engaged towards the same target event.

**Figure 6.**
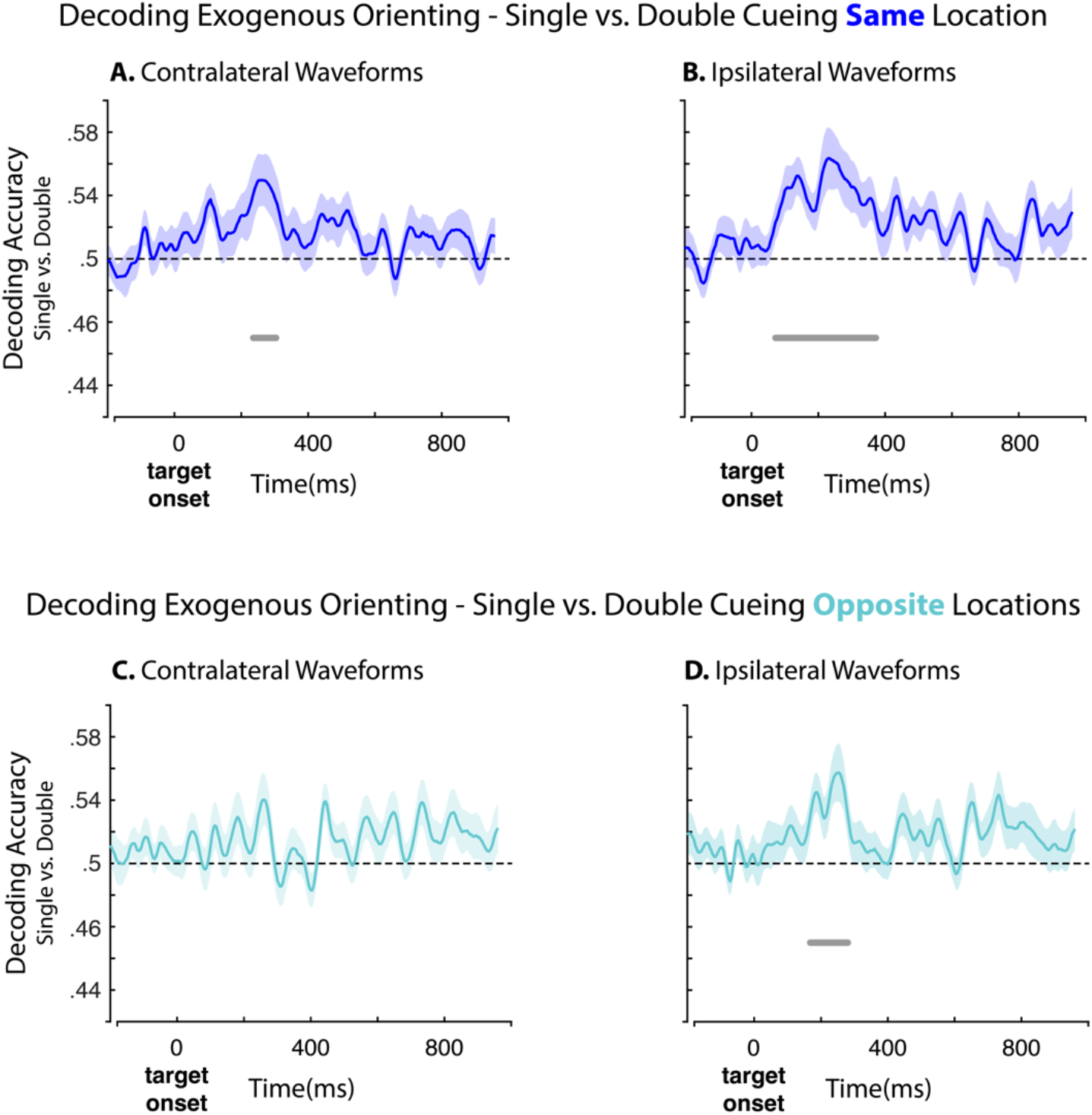
Multivariate classification of cueing conditions based on target-related ERPs for contra- and ipsilateral channels (i.e., cue valid minus cue invalid. We assessed classification performance for contralateral trials to the target when both cues indicated the same location (**A**) and the opposite location (**C**). We repeated this analysis for ipsilateral trials (**C**) and (**D**). Grey lines indicate significant clusters against chance-level based on cluster-corrected t-tests. Bold lines represent the averaged time series and shaded areas of the SEM.

Lastly, we decoded cue conditions from target-related LAI. Note that LAI was computed relative to the target’s location. For this analysis, we decoded cue conditions based on cue validity effects: We subtracted trials when the cue was valid versus invalid (see Methods for details). We opted to decode cueing conditions from single cueing versus double-cueing opposite location trials only. This analysis revealed a significant cluster occurring from −84 ms to 284 ms relative to target onset (Figure 7A). We further examined the weights of the model and observed that above chance decoding was related to modulations of LAI occurring over posterior regions (Figure 7B).

**Figure 7.**
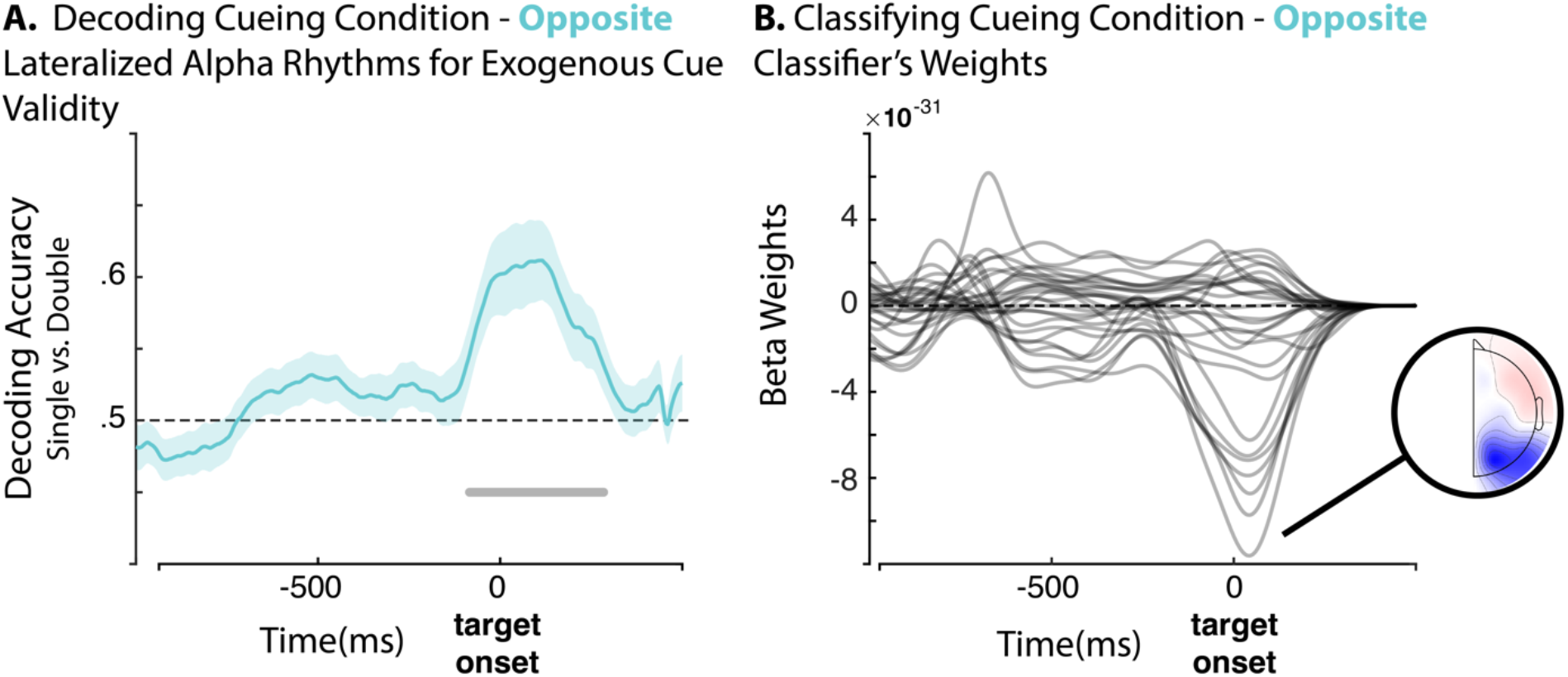
Multivariate classification of cueing conditions (i.e., single versus double cueing for opposite cueing locations) based on target-related lateralized alpha power cue validity effect (i.e., cue valid minus cue invalid). We assessed classification performance for decoding cueing condition (**A**) against chance-level. Grey lines indicate significant clusters against chance-level based on cluster-corrected t-tests. Bold lines represent the averaged time series and shaded areas of the SEM. The corresponding weights of the classifiers across the time series (**B**), and the topography for the minimal value occurring 44ms after target onset.

## Discussion

Exogenous and endogenous orienting correspond to distinct control systems of visuospatial attention (Chica et al., 2013). And yet, previous studies show that these attention processes interact with each other under certain conditions, thereby challenging the idea that they are fully separable (Ahrens et al., 2019; Grubb et al., 2015; Hopfinger & West, 2006; MacLean et al., 2009). Berger et al. (2005) have argued that these interactions indicate that exogenous and endogenous attention are capable of mutual interference – a viewpoint that implies limited attention resources between them. The present work explored this perspective by examining the influence of endogenous attention on exogenous attention processes, and then examined whether this influence involves alpha-band activity.

We first confirmed that endogenous attention modulates cue-related and target-related processes of exogenous attention, while we show that these effects varied according to the direction of endogenous attention relative to that of exogenous attention. Thus, our findings confirmed that exogenous attention is not immune to top-down processes involved in endogenous attention (Landry et al., 2021; Pinto et al., 2013). Specifically, previous reports show that peripheral visual cues yield early (>200 ms) and late (<200 ms) neurophysiological brain activity related to exogenous attention (Martín-Arévalo et al., 2014; Störmer et al., 2019). The present work reveals that the concurrent engagement of endogenous attention alters the processing of the exogenous cue early on (Figure 2). This early influence implies that interactions between exogenous and endogenous attention processes occur at lower levels of sensory processing, which is consistent with neuroimaging assays showing that both attention processes involve modulations of the primary visual cortex (Dugué et al., 2020; Müller & Ebeling, 2008). Moreover, examining the weights of the SVM model shows that the processing of the exogenous cue encompasses a lateralized posterior topographical pattern that maps onto contra- and ipsi-lateral neural responses relative to the location of the exogenous cue (Figure 4). Whenever endogenous attention was engaged to the opposite location, we observed modulations of this lateralized component between 200 ms and 600 ms (Figure 4). The influence of endogenous attention cue-related processing is therefore not limited to early processes, but also involves further stages of processing. Together, these results entail that the concurrent engagement of endogenous attention alters the processing of the peripheral cue in a significant manner across multiple stages of processing along the visual pathway.

We also observed that endogenous attention interferes with the exogenous cue validity effect, both at the behavioral and neurophysiological levels. This outcome entails that, in addition to altering the processing of the peripheral cue, endogenous attention likewise interferes with exogenous attention’s processing of the target event. Our results align with a previous report showing that exogenous and endogenous interact early during visual processing of target stimuli – an effect occurring around the N1 target-related ERP component (Hopfinger & West, 2006). Our work extends these findings by showing that the impact of endogenous attention on the exogenous cue validity effect involves both ipsi- and contra-lateral neural responses around 200 ms after target onset (Figure 6). Importantly, we observed this effect both when endogenous and exogenous attention are engaged towards the same location and when they are engaged toward opposite locations. Hence, the mere engagement of endogenous attention processes, regardless of its direction, alters the cueing effects of exogenous attention. This outcome is consistent with the idea that exogenous and endogenous processes share neural processing resources at the sensory level, while this overlap leads to interference during target processing.

Lastly, having confirmed that endogenous attention alters cue-related and target-related exogenous attention, we evaluated our main hypothesis and tested whether these effects involve the power and phase of alpha band activity. Our hypothesis follows from past work that highlights the centrality of alpha power and phase across both exogenous and endogenous attention processes (D’Andrea et al., 2019; e.g., Keefe & Störmer, 2021; Störmer et al., 2016; Thut et al., 2006). We accordingly hypothesized that both attention processes would tap into the same neural mechanisms of visuospatial attention, thereby leading to interactions between them (Busse et al., 2008). To test our hypothesis, we trained classifiers to decode the direction of exogenous attention based on the instantaneous power and phase of alpha rhythms in the single cueing condition and then tested them in context of the double cueing condition. These classifiers were therefore naive to the effects of endogenous attention. Our results show that the direction of endogenous attention during double cueing determined classification performances: The classifiers performed above chance level when endogenous attention was oriented towards the same location as exogenous attention and below it whenever endogenous attention was oriented towards the opposite location (Figure 3). This pattern occurred for both the power and phase of alpha rhythms. The strong influence of endogenous attention over classifiers trained to decode exogenous attention supports the idea of shared neurophysiological processes across the power and phase of alpha rhythms between them. Furthermore, consistent with our behavioral results, we show that concurrent engagement of endogenous attention to the opposite location alters lateralized alpha activity pertaining to exogenous cue validity, and that this effect corresponds to a posterior topographical pattern (Figure 7). Therefore, our findings reveal that alpha-band activity captures the interference of endogenous attention with the cueing benefit of exogenous attention.

According to prevailing views, synchronized activity in the range of alpha rhythms at the sensory level reflects inhibitory and excitatory mechanisms during selective attention (Bacigalupo & Luck, 2019). Different pieces of evidence support this viewpoint. First and foremost, visuospatial attention links with desynchronized alpha rhythms in the contralateral posterior sensory region and synchronized alpha rhythms in the ipsilateral posterior region. This lateralized pattern of alpha band activity is thought to reflect increased neuronal excitability for the processing of attended sensory events and increased neuronal inhibition for the suppression of unattended inputs (Sauseng et al., 2005; Wöstmann et al., 2019). Second, fluctuations of ongoing alpha rhythms have been linked to various aspects related to sensory processing, such as task performance (e.g., Bollimunta et al., 2008), visual perception (e.g., van Dijk et al., 2008), and perceptual representations (e.g., Zhou et al., 2021). Lastly, various studies relate changes in alpha-band activity to states of neuronal excitability (Dougherty et al., 2017; Goldman et al., 2002; Iemi et al., 2022). We accordingly hypothesized that both exogenous and endogenous would yield perceptual benefits via the same neural mechanisms at the sensory level, as indexed by change in alpha-band activity. This hypothesis follows from the idea that the relationship between endogenous and exogenous attention is marked by competition for limited perceptual resources, which yields mutual interference between them (Berger et al., 2005). Our results therefore align with this viewpoint by showing that both attention processes tap into the same mechanisms for the facilitation of sensory processing and the suppression of irrelevant inputs (i.e., modulations of alpha-band brain activity), which yields interference between them. Specifically, we can appreciate how this overlap can lead to conflicting patterns of neural excitability and inhibition at the sensory level whenever exogenous and endogenous attention engage opposite sides of the visual field. In sum, interference of endogenous attention over exogenous attention processes occurs, at least in part, because both systems share the same neural processes at the sensory level.

Exogenous and endogenous represent independent attention systems capable of mutual interference. Exploring whether such interference occurs via alpha-band neurophysiological activity, we tested the influence of endogenous attention on exogenous attention processes and found evidence that they share the same neurophysiological resources to modulate neuronal excitability and inhibition at the sensory level. Therefore, while both systems involve distinct modes of control and separable neural processes, their effects on exogenous and endogenous attention overlap at the sensory level of processing.

## Acknowledgments

M.L. acknowledges support from the Fonds de Recherche du Québec – Nature et Technologies. J.D.C. acknowledges a doctoral fellowship from the Natural Science and Engineering Research Council of Canada. S.B. is supported by the NIH (R01 EB026299), a Discovery grant from the Natural Science and Engineering Research Council of Canada (436355-13), the CIHR Canada research Chair in Neural Dynamics of Brain Systems, the Brain Canada Foundation with support from Health Canada, and the Innovative Ideas program from the Canada First Research Excellence Fund, awarded to McGill University for the Healthy Brains for Healthy Lives initiative (1c-II-14). This work was supported by a grant from CIHR and Canada Research Chair to A.R. J.S. acknowledges grants from the ANR (France): ANR-16-ASTR-0014 and ANR-17-EURE-0017. We would like to thank Anna Prokusheva for her help collecting the data. Lastly, we would also like to thank Barry Giesbrecht, Mathieu Roy, Dave Saint-Amour, Vincent de Gardelle, and Viola Störmer for helpful comments about this research project.

## Contributions

M.L. and A.R. discussed and dreamed up the study. M.L. and J.D.C. designed experiments, collected data, and performed analyses. J.S., S.B. and A.R. provided feedback throughout the project. All authors wrote the manuscript. A.R secured funding and provided lab space and equipment to pursue this study.

## Data and Codes availability

Data and codes will be available on OSF: https://osf.io/z8wkp/?view_only=9cc6c28dc5504ec48bf933f002d6ee8c

## References

Ahrens, M.-M., Veniero, D., Freund, I. M., Harvey, M., & Thut, G. (2019, 2019/08/01/). Both dorsal and ventral attention network nodes are implicated in exogenously driven visuospatial anticipation. Cortex, 117, 168–181.

Bacigalupo, F., & Luck, S. J. (2019). Lateralized suppression of alpha-band EEG activity as a mechanism of target processing. Journal of Neuroscience, 39(5), 900–917.

Bae, G.-Y., & Luck, S. J. (2018). Dissociable decoding of spatial attention and working memory from EEG oscillations and sustained potentials. Journal of Neuroscience, 38(2), 409–422.

Bagherzadeh, Y., Baldauf, D., Pantazis, D., & Desimone, R. (2020). Alpha synchrony and the neurofeedback control of spatial attention. Neuron, 105(3), 577–587.

Berger, A., Henik, A., & Rafal, R. (2005). Competition between endogenous and exogenous orienting of visual attention. Journal of experimental psychology: General, 134(2), 207.

Bollimunta, A., Chen, Y., Schroeder, C. E., & Ding, M. (2008). Neuronal Mechanisms of Cortical Alpha Oscillations in Awake-Behaving Macaques. The Journal of Neuroscience, 28(40), 9976.

Bowling, J. T., Friston, K. J., & Hopfinger, J. B. (2020). Top-down versus bottom-up attention differentially modulate frontal–parietal connectivity. Human brain mapping, 41(4), 928–942.

Brainard, D. H., & Vision, S. (1997). The psychophysics toolbox. Spatial vision, 10, 433–436.

Buschman, T. J., & Miller, E. K. (2007). Top-down versus bottom-up control of attention in the prefrontal and posterior parietal cortices. science, 315(5820), 1860–1862.

Busse, L., Katzner, S., & Treue, S. (2008). Temporal dynamics of neuronal modulation during exogenous and endogenous shifts of visual attention in macaque area MT. Proceedings of the national academy of sciences.

Carrasco, M. (2011). Visual attention: The past 25 years. Vision research, 51(13), 1484–1525.

Chica, A. B., Bartolomeo, P., & Lupiáñez, J. (2013). Two cognitive and neural systems for endogenous and exogenous spatial attention. Behavioural brain research, 237, 107–123.

Chica, A. B., Martín-Arévalo, E., Botta, F., & Lupiánez, J. (2014). The Spatial Orienting paradigm: How to design and interpret spatial attention experiments. Neuroscience and Biobehavioral Reviews, 40, 35–51.

Cohen, M. X. (2014). Analyzing neural time series data: theory and practice. MIT Press.

Corbetta, M., Patel, G., & Shulman, G. L. (2008). The reorienting system of the human brain: from environment to theory of mind. Neuron, 58(3), 306–324.

Corbetta, M., & Shulman, G. L. (2002). Control of goal-directed and stimulus-driven attention in the brain. Nature reviews neuroscience, 3(3), 201–215.

D’Andrea, A., Chella, F., Marshall, T. R., Pizzella, V., Romani, G. L., Jensen, O., & Marzetti, L. (2019). Alpha and alpha-beta phase synchronization mediate the recruitment of the visuospatial attention network through the Superior Longitudinal Fasciculus. NeuroImage, 188, 722–732.

Dougherty, K., Cox, M. A., Ninomiya, T., Leopold, D. A., & Maier, A. (2017). Ongoing Alpha Activity in V1 Regulates Visually Driven Spiking Responses. Cerebral Cortex, 27(2), 1113–1124.

Dugué, L., Merriam, E. P., Heeger, D. J., & Carrasco, M. (2020, 2020/12/04). Differential impact of endogenous and exogenous attention on activity in human visual cortex. Scientific Reports, 10(1), 21274.

Gelman, A., & Hill, J. (2006). Data Analysis Using Regression and Multilevel/Hierarchical Models. Cambridge University Press.

Goldman, R. I., Stern, J. M., Engel, J., Jr., & Cohen, M. S. (2002). Simultaneous EEG and fMRI of the alpha rhythm. Neuroreport, 13(18).

Grubb, M. A., White, A. L., Heeger, D. J., & Carrasco, M. (2015). Interactions between voluntary and involuntary attention modulate the quality and temporal dynamics of visual processing. Psychonomic bulletin & review, 22(2), 437–444.

Haufe, S., Meinecke, F., Görgen, K., Dähne, S., Haynes, J.-D., Blankertz, B., & Bießmann, F. (2014). On the interpretation of weight vectors of linear models in multivariate neuroimaging. NeuroImage, 87, 96–110.

Hong, X., Bo, K., Meyyappan, S., Tong, S., & Ding, M. (2020). Decoding attention control and selection in visual spatial attention. Human brain mapping, 41(14), 3900–3921.

Hopfinger, J. B., & West, V. M. (2006). Interactions between endogenous and exogenous attention on cortical visual processing. NeuroImage, 31(2), 774–789.

Iemi, L., Gwilliams, L., Samaha, J., Auksztulewicz, R., Cycowicz, Y. M., King, J.-R., Nikulin, V. V., Thesen, T., Doyle, W., Devinsky, O., Schroeder, C. E., Melloni, L., & Haegens, S. (2022, 2022/02/15/). Ongoing neural oscillations influence behavior and sensory representations by suppressing neuronal excitability. NeuroImage, 247, 118746.

Jonides, J. (1981). Voluntary versus automatic control over the mind’s eye’s movement. Attention and performance IX, 9, 187–203.

Keefe, J. M., & Störmer, V. S. (2021, 2021/01/15/). Lateralized alpha activity and slow potential shifts over visual cortex track the time course of both endogenous and exogenous orienting of attention. NeuroImage, 225, 117495.

Kleiner, M., Brainard, D., Pelli, D., Ingling, A., Murray, R., & Broussard, C. (2007). What’s new in Psychtoolbox-3? [ECVP Abstract Supplement]. Perception, 36(14).

Klimesch, W. (2012, 2012/12/01/). Alpha-band oscillations, attention, and controlled access to stored information. Trends in cognitive sciences, 16(12), 606–617.

Landry, M., Da Silva Castanheira, J., Sackur, J., & Raz, A. (2021). Investigating how the modularity of visuospatial attention shapes conscious perception using type I and type II signal detection theory. Journal of Experimental Psychology: Human Perception and Performance, 47(3), 402.

Luck, S. J., Hillyard, S. A., Mouloua, M., & Hawkins, H. L. (1996). Mechanisms of visual–spatial attention: Resource allocation or uncertainty reduction? Journal of Experimental Psychology: Human Perception and Performance, 22(3), 725.

MacLean, K. A., Aichele, S. R., Bridwell, D. A., Mangun, G. R., Wojciulik, E., & Saron, C. D. (2009). Interactions between endogenous and exogenous attention during vigilance. Attention, Perception, & Psychophysics, 71(5), 1042–1058.

Martín-Arévalo, E., Chica, A. B., & Lupiáñez, J. (2014, 2014/03/01/). Electrophysiological modulations of exogenous attention by intervening events. Brain and Cognition, 85, 239–250.

Mathewson, K. E., Gratton, G., Fabiani, M., Beck, D. M., & Ro, T. (2009). To see or not to see: prestimulus α phase predicts visual awareness. Journal of Neuroscience, 29(9), 2725–2732.

Müller, H. J., & Rabbitt, P. M. A. (1989). Spatial cueing and the relation between the accuracy of “where” and “what” decisions in visual search. The Quarterly Journal of Experimental Psychology, 41(4), 747–773.

Müller, N. G., & Ebeling, D. (2008, 2008/04/01/). Attention-modulated activity in visual cortex—More than a simple ‘spotlight’. NeuroImage, 40(2), 818–827.

Nobre, A. C., Sebestyen, G. N., Gitelman, D. R., Mesulam, M. M., Frackowiak, R. S., & Frith, C. D. (1997). Functional localization of the system for visuospatial attention using positron emission tomography. Brain, 120(3), 515–533.

Peelen, M. V., Heslenfeld, D. J., & Theeuwes, J. (2004, 2004/06/01/). Endogenous and exogenous attention shifts are mediated by the same large-scale neural network. NeuroImage, 22(2), 822–830.

Pelli, D. G. (1997). The VideoToolbox software for visual psychophysics: Transforming numbers into movies. Spatial vision, 10(4), 437–442.

Peylo, C., Hilla, Y., & Sauseng, P. (2021, 2021/09/01/). Cause or consequence? Alpha oscillations in visuospatial attention. Trends in Neurosciences, 44(9), 705–713.

Pinto, Y., van der Leij, A. R., Sligte, I. G., Lamme, V. A. F., & Scholte, H. S. (2013). Bottom-up and top-down attention are independent. Journal of Vision, 13(3), 16–16.

Posner, M. I. (1980). Orienting of attention. Quarterly journal of experimental psychology, 32(1), 3–25.

Posner, M. I., Snyder, C. R., & Davidson, B. J. (1980). Attention and the detection of signals. Journal of experimental psychology: General, 109(2), 160.

Ristic, J., & Kingstone, A. (2012). A new form of human spatial attention: automated symbolic orienting. Visual Cognition, 20(3), 244–264.

Ristic, J., & Landry, M. (2015). Combining attention: a novel way of conceptualizing the links between attention, sensory processing, and behavior. Attention, Perception, & Psychophysics, 77(1), 36–49.

Santangelo, V., & Spence, C. (2008). Is the exogenous orienting of spatial attention truly automatic? Evidence from unimodal and multisensory studies. Consciousness and cognition, 17(3), 989–1015.

Sauseng, P., Klimesch, W., Stadler, W., Schabus, M., Doppelmayr, M., Hanslmayr, S., Gruber, W. R., & Birbaumer, N. (2005). A shift of visual spatial attention is selectively associated with human EEG alpha activity. European Journal of Neuroscience, 22(11), 2917–2926.

Störmer, V. S., Feng, W., Martinez, A., McDonald, J. J., & Hillyard, S. A. (2016). Salient, irrelevant sounds reflexively induce alpha rhythm desynchronization in parallel with slow potential shifts in visual cortex. Journal of Cognitive Neuroscience, 28(3), 433–445.

Störmer, V. S., McDonald, J. J., & Hillyard, S. A. (2019). Involuntary orienting of attention to sight or sound relies on similar neural biasing mechanisms in early visual processing. Neuropsychologia, 132, 107122.

Tadel, F., Baillet, S., Mosher, J. C., Pantazis, D., & Leahy, R. M. (2011). Brainstorm: a user-friendly application for MEG/EEG analysis. Computational intelligence and neuroscience, 2011, 8.

Tallon-Baudry, C., Bertrand, O., Delpuech, C., & Pernier, J. (1996). Stimulus specificity of phase-locked and non-phase-locked 40 Hz visual responses in human. Journal of Neuroscience, 16(13), 4240–4249.

Thut, G., Nietzel, A., Brandt, S. A., & Pascual-Leone, A. (2006). α-Band electroencephalographic activity over occipital cortex indexes visuospatial attention bias and predicts visual target detection. Journal of Neuroscience, 26(37), 9494–9502.

van Dijk, H., Schoffelen, J.-M., Oostenveld, R., & Jensen, O. (2008). Prestimulus Oscillatory Activity in the Alpha Band Predicts Visual Discrimination Ability. The Journal of Neuroscience, 28(8), 1816.

Wagenmakers, E.-J. (2007). A practical solution to the pervasive problems of p values. Psychonomic bulletin & review, 14(5), 779–804.

Woldorff, M. G. (1993). Distortion of ERP averages due to overlap from temporally adjacent ERPs: Analysis and correction. Psychophysiology, 30(1), 98–119.

Wöstmann, M., Alavash, M., & Obleser, J. (2019). Alpha oscillations in the human brain implement distractor suppression independent of target selection. Journal of Neuroscience, 39(49), 9797–9805.

Wright, R. D., & Ward, L. M. (2008). Orienting of attention. Oxford University Press.

Zhou, Y. J., Iemi, L., Schoffelen, J.-M., de Lange, F. P., & Haegens, S. (2021). Alpha oscillations shape sensory representation and perceptual sensitivity. Journal of Neuroscience, 41(46), 9581–9592.

